# NAP1L1 and NAP1L4 binding to Hypervariable Domain of Chikungunya Virus nsP3 Protein is bivalent and requires phosphorylation

**DOI:** 10.1101/2021.05.19.444900

**Authors:** Francisco Dominguez, Nikita Shiliaev, Tetyana Lukash, Peter Agback, Oksana Palchevska, Joseph R. Gould, Chetan D. Meshram, Peter E. Prevelige, Todd J. Green, Tatiana Agback, Elena I. Frolova, Ilya Frolov

## Abstract

Chikungunya virus (CHIKV) is one of the most pathogenic members of the *Alphavirus* genus in the *Togaviridae* family. Within the last two decades, CHIKV has expanded its presence to both hemispheres and is currently circulating in both Old and New Worlds. Despite the severity and persistence of the arthritis it causes in humans, no approved vaccines or therapeutic means have been developed for CHIKV infection. Replication of alphaviruses, including CHIKV, is determined not only by their nonstructural proteins, but also by a wide range of host factors, which are indispensable components of viral replication complexes (vRCs). Alphavirus nsP3s contain hypervariable domains (HVDs), which encode multiple motifs that drive recruitment of cell- and virus-specific host proteins into vRCs. Our previous data suggested that NAP1 family members are a group of host factors that may interact with CHIKV nsP3 HVD. In this study, we performed a detailed investigation of the NAP1 function in CHIKV replication in vertebrate cells. Our data demonstrate that i) the NAP1-HVD interactions have strong stimulatory effects on CHIKV replication; ii) both NAP1L1 and NAP1L4 interact with the CHIKV HVD; iii) NAP1 family members interact with two motifs, which are located upstream and downstream of the G3BP-binding motifs of CHIKV HVD; iv) NAP1 proteins interact only with a phosphorylated form of CHIKV HVD and HVD phosphorylation is mediated by CK2 kinase; v) NAP1 and other families of host factors redundantly promote CHIKV replication and their bindings have additive stimulatory effects on viral replication.

**IMPORTANCE:** Cellular proteins play critical roles in the assembly of alphavirus replication complexes (vRCs). Their recruitment is determined by the viral nonstructural protein 3 (nsP3). This protein contains a long, disordered hypervariable domain (HVD), which encodes virus-specific combinations of short linear motifs interacting with host factors during vRC assembly. Our study defined the binding mechanism of NAP1 family members to CHIKV HVD and demonstrated a stimulatory effect of this interaction on viral replication. We showed that interaction with NAP1L1 is mediated by two HVD motifs and requires phosphorylation of HVD by CK2 kinase. Based on the accumulated data, we present a map of the binding motifs of the critical host factors currently known to interact with CHIKV HVD. It can be used to manipulate cell specificity of viral replication and pathogenesis, and to develop a new generation of vaccine candidates.

## INTRODUCTION

Members of the *Alphavirus* genus in the *Togaviridae* family are circulating on all continents (1). Most of them are transmitted by mosquito vectors between vertebrate hosts, in which they cause diseases of different severities. Based on their geographical distribution, alphaviruses are referred to as the New World (NW) and the Old World (OW) alphaviruses. However, recently, chikungunya virus (CHIKV) became an exception, as within the last two decades, it has dramatically expanded its circulation area and is currently present in both the New and Old Worlds (2, 3). In humans, CHIKV causes a highly debilitating disease characterized by rash, fever and, most importantly, severe joint pain that can persist for years (4). Thus, CHIKV represents an unquestionable public health threat, but to date, no licensed vaccines or therapeutic means have been developed for this infection. As for other alphaviruses, the molecular mechanisms of CHIKV replication, virus-host interactions and pathogenesis remain to be better understood.

CHIKV genome is represented by a single-stranded RNA (G RNA) of positive polarity (5–7). Similar to cellular mRNAs, it contains a cap and a poly(A) tail at the 5’ and 3’ termini, respectively. Upon delivery into the cytoplasm by viral particles or RNA transfection, the genome is translated into nonstructural (ns) polyproteins P123 and P1234. They are targeted to the plasma membrane and sequentially processed into individual viral nonstructural proteins (nsP1-to-4) by the nsP2-encoded protease (8). nsPs serve as viral components of the replication complexes (vRCs) that synthesize dsRNA replication intermediates on the G RNA template, and new G and subgenomic (SG) RNAs. Processing of ns polyproteins regulates the template specificities of vRCs at different steps of viral replication. The partially processed P123+nsP4 complex functions in the negative strand RNA synthesis, and the completely processed nsPs mediate synthesis of the positive strand G and SG RNAs, but cannot synthesize the negative strand RNAs.

The enzymatic functions of nsP1, nsP2 and nsP4 in the synthesis of virus-specific RNAs and in the unique cascade of capping reactions are relatively well understood (7, 9). However, to date no direct enzymatic activities in G RNA replication and transcription of SG RNA have been ascribed for nsP3 protein. Nevertheless, nsP3 is indispensable for viral RNA synthesis, and mutations in this protein have deleterious effects on viral replication (10–13). nsP3 contains two N-terminal, conserved, structured domains [macro domain and alphavirus unique domain (AUD)], whose functions remain to be determined, and the C-terminal hypervariable domain (HVD) (14–16). Our previous studies and those of other groups demonstrated that alphavirus HVDs encode a variety of short motifs, which recruit specific combinations of host factors into viral vRCs, and alterations of HVD interactions with host proteins either have strong negative effects on the rates of viral replication or can make alphaviruses nonviable (13, 17–23).

The CHIKV nsP3 HVD is intrinsically disordered and was shown to interact with cell-specific sets of host proteins (17, 19, 21, 24, 25). The accumulated data about its interactions with host factors can be summarized as follows. i) G3BP1 and G3BP2 are the critical host factors for CHIKV replication in vertebrate cells (17, 26). In mosquito cells, the insect-specific homolog RIN1 appears to have the same pro-viral function as G3BPs of vertebrates (27). Mutations in G3BP-binding sites of nsP3 HVD or knock out (KO) of both G3BP1/2 genes in mammalian cells make CHIKV not viable (17). ii) Binding of only G3BPs to CHIKV HVD is insufficient for vRC formation and function, and the variants that have only G3BP-binding sites in their HVDs remain nonviable (19). Interactions with other families of host proteins have stimulatory effects on viral replication. These additional interacting partners include the SH3 domain-containing proteins (BIN1, CD2AP and SH3KBP1), members of the FHL family (FHL1-to-3) and NAP1 (nucleosome assembly protein 1) family members (19, 23). However, this list probably remains incomplete. iii) The distinguishing characteristic of the HVD-binding factors is the high redundancy of their function. It allows viral CHIKV to utilize multiple members of the families and different protein families to facilitate vRC formation and function. The use of more than one family member likely promotes viral replication in a wider range of tissues and host species, in which the presence of particular protein factors may vary. iv) We have recently characterized by NMR spectroscopy and genetic approaches the binding sites of the members of G3BP and FHL families, and the SH3 domain-containing proteins. The additive pro-viral effects of these interactions were demonstrated in biological studies (23, 24). v) Binding of NAP1 proteins to CHIKV HVD and its biological significance remains poorly explored, although this interaction appears to be CHIKV-specific and is required for efficient viral replication in vertebrate cells. NAP1 proteins are not extensively characterized, but are known to be essential for cell growth. NAP1L1 is also believed to be involved in DNA replication with higher expression in rapidly proliferating cells and in tumors. NAP1L1 and NAP1L4 were also implicated in regulation of cellular transcription (28–30).

In this new study, we used a wide variety of approaches to identify the binding sites of NAP1L1 and NAP1L4 in CHIKV HVD and demonstrated the pro-viral role of this interaction in CHIKV replication. Our data reveal a unique mechanism of NAP1L1 binding to HVD. It is determined by two HVD-encoded motifs located upstream and downstream of the G3BP-binding sites. However, NAP1-HVD interaction remains G3BP-independent. Importantly, interaction of NAP1 proteins with CHIKV nsP3 is dependent on HVD phosphorylation by CK2 kinase The presence of only NAP1-binding sites in CHIKV HVD in addition to the G3BP-binding motifs is sufficient for CHIKV viability, but binding of other host factors has strong stimulatory effect on viral replication. These results additionally support our overall hypothesis regarding redundant functions of alphavirus HVD-interacting partners.

## RESULTS

### NAP1L1 interacts with the C-terminal fragment of CHIKV nsP3 HVD both in the presence and absence of G3BP

In previous studies, we and other groups have identified sets of host factors that interact with CHIKV nsP3 HVD in human and mouse cells (18–23). These HVD-interacting partners include the members of G3BP family (G3BP1 and G3BP2), the SH3 domain-containing proteins (CD2AP, BIN1 and SH3KBP1), FHL family members (FHL1, FHL2 and FHL3) and proteins of the NAP1 family (NAP1L1 and NAP1L4). To further understand the mechanism of NAP1L1 and NAP1L4 interactions with CHIKV HVD and their roles in viral replication, we first confirmed that in CHIKV-infected cells, NAP1L1 and NAP1L4 are present in the nsP3 complexes. NIH 3T3 cells were infected with CHIKV 181/25 and stained with nsP3-, NAP1L1- and NAP1L4-specific Abs. As demonstrated in Fig. 1, both NAP1 proteins clearly accumulated in large nsP3-containing cytoplasmic complexes.

**FIG 1.**
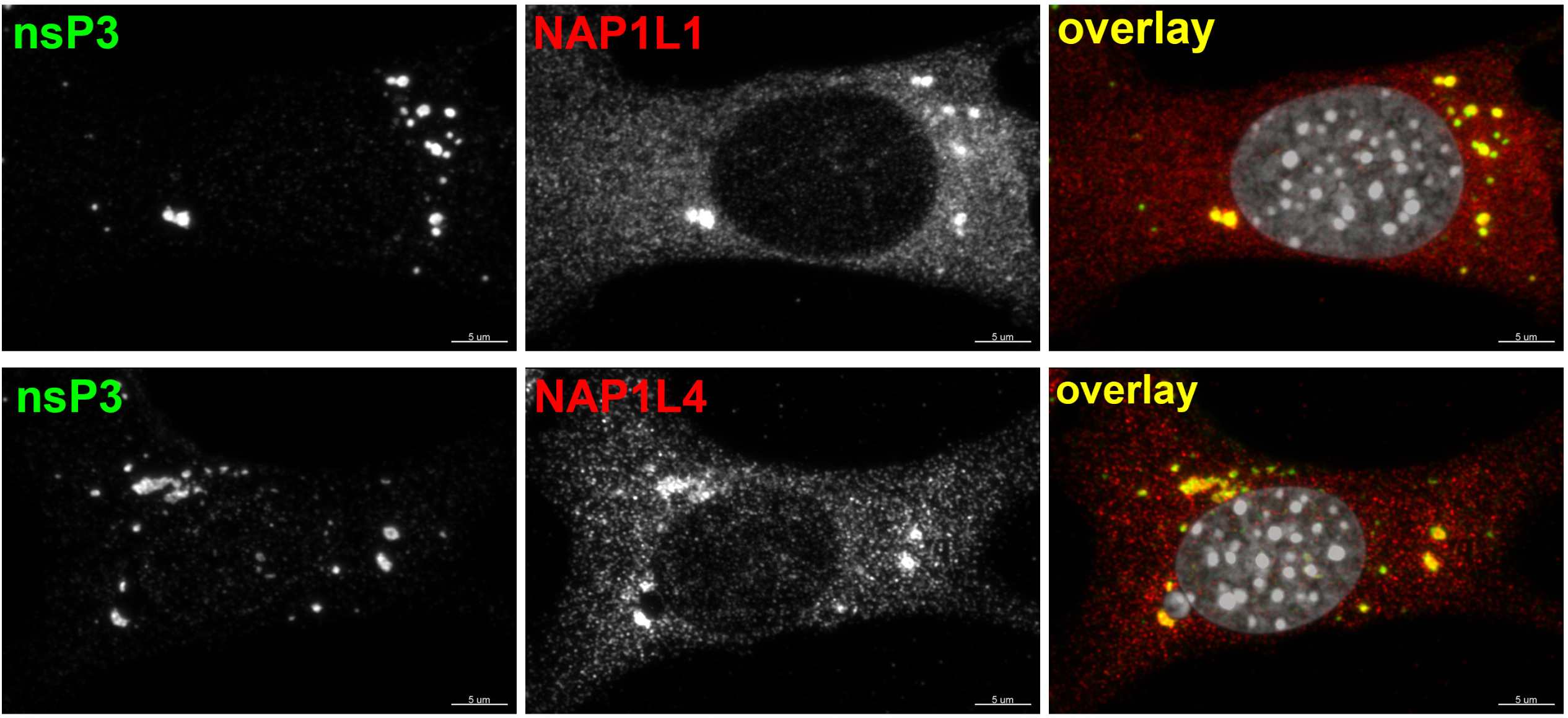
NAP1L1 and NAP1L4 accumulate in CHIKV nsP3 complexes. NIH 3T3 cells were infected with CHIKV 181/25 at an MOI of 10 PFU/cell in 8-well Ibidi plates. At 7 h p.i., cells were fixed and stained with CHIKV nsP3- and NAP1-specific Abs, and DAPI as described in Materials and Methods. Images were acquired on a Zeiss LSM 800 confocal microscope in Airyscan mode with a 63X 1.4NA PlanApochromat oil objective.

Next, we intended to identify the NAP1L1-binding motif in CHIKV nsP3 HVD. As in our recent studies (19), we utilized a set of Venezuelan equine encephalitis virus (VEEV) replicons (VEErep) to express different combinations of CHIKV HVD fragments fused with Flag-GFP (Fig. 2A). The HVD was divided into 4 fragments either having wild type (wt) amino acid (aa) sequence (fragments A, B, C and D) or having aa sequences randomized by changing the positions of closely located aa (fragments 1, 2 and 3). These modifications were aimed at eliminating the known and other potential binding sites of host proteins in the individual fragments. In the (3C) fragment, only the C-terminal 16-aa-long peptide of fragment C was left intact, and the remaining two thirds of the fragment were randomized. The designed replicons were packaged into infectious viral particles using VEEV TC-83-based helper RNA (see Materials and Methods for details). Then, NIH 3T3 cells and their *G3bp* dKO derivatives (17) were infected with packaged replicons, and protein complexes were isolated using magnetic beads loaded with Flag-specific MAbs. Only the ABCD-, 12CD- and 12(3C)D-encoding constructs demonstrated the ability to efficiently co-immunoprecipitate NAP1L1 and NAP1L4 (Fig. 2B). Thus, interaction of NAP1 family members with CHIKV HVD was determined by the C-terminal HVD fragment (3C)D, which contained binding sites of the G3BP1/2 (fragment D), and the last 16 aa of the upstream fragment C. Since the results generated on *G3bp* dKO and parental NIH 3T3 cells were similar (Fig. 2B), we concluded that both NAP1 proteins can bind to CHIKV HVD in the absence of the G3BP interaction with CHIKV HVD. However, since G3BP- and NAP1-binding sites are so closely located, the possibility of cooperativity in interactions of these two protein families with CHIKV HVD could not be completely ruled out.

**FIG. 2.**
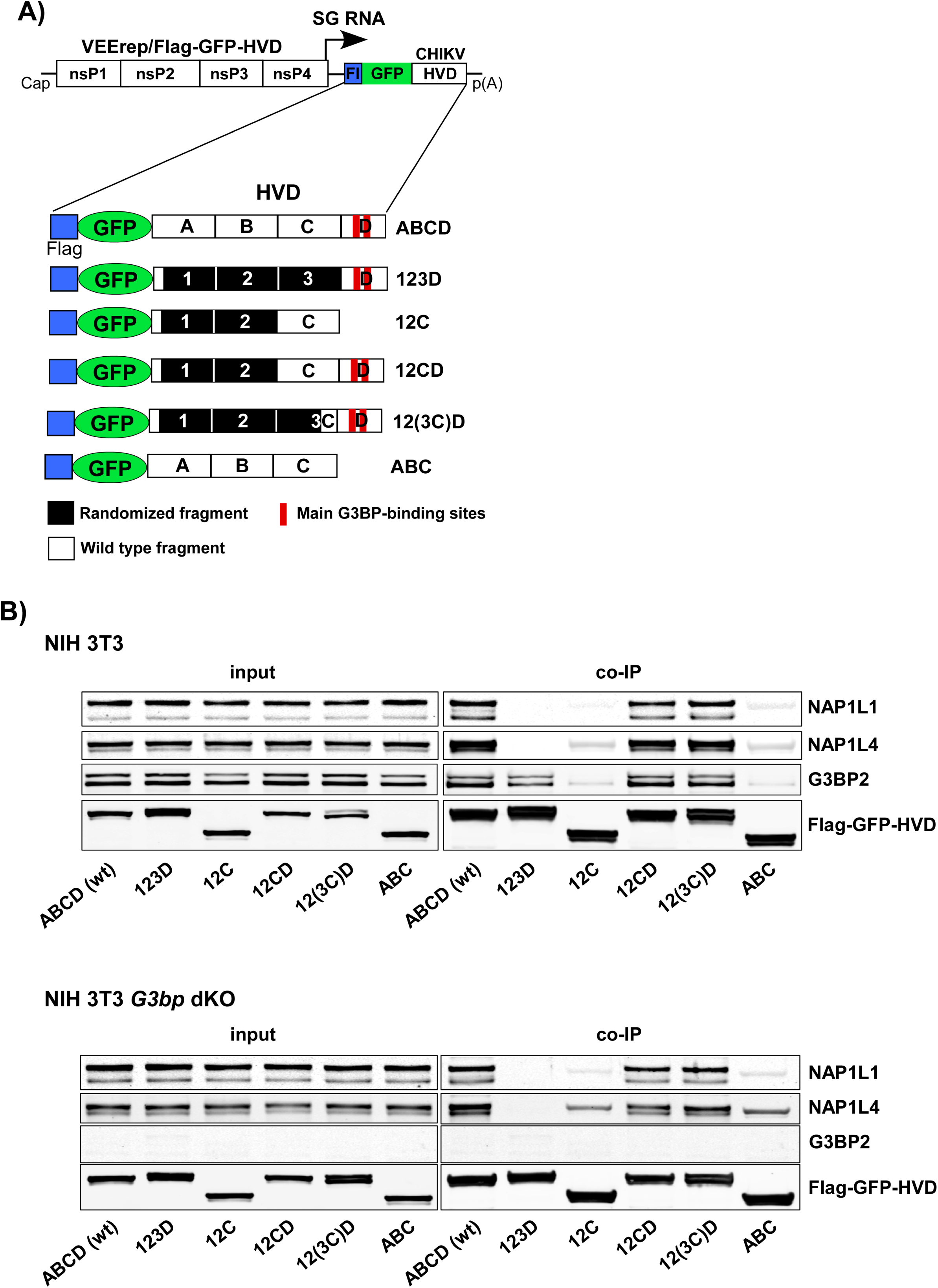
NAP1 family members are capable of interacting with the C-terminal fragment of CHIKV HVD independently of G3BP. (A) The schematic presentation of VEEV replicons encoding different variants of CHIKV HVD fused with Flag-GFP. A, B, C and D indicate wt aa sequences. Fragments 1, 2, and 3 have the aa sequences of the corresponding A, B and C fragments randomized. In fragment (3C) only the C-terminal 16-aa-long peptide had the wt sequence. Wt and mutated fragments are additionally indicated by open and black boxes, respectively. (B) NIH 3T3 cells and their *G3bp* dKO derivatives were infected at an MOI of 10 inf.u/cell with the packaged VEEV replicons expressing the indicated fragments of CHIKV HVD. Cells were collected at 4 h p.i. Protein complexes were isolated on magnetic beads as described in Materials and Methods and analyzed by Western blot using NAP1L1-, NAP1L4-, G3BP2 and Flag-specific Abs and corresponding secondary Abs. Membranes were scanned on a Li-Cor imager. The experiment was repeated three times with similar results.

### Mutations in (3C) fragment of CHIKV nsP3 HVD affect viral replication

Next, we further characterized function of the identified (3C) fragment in CHIKV replication. For this analysis, we applied a CHIKV 181/25 variant encoding 123(3C)D HVD, termed CHIKV/(3C)/GFP (Fig. 3A). nsP3 HVD of this virus contained no binding motifs of the FHL or the SH3 domain-containing proteins. Its replication was expected to rely only on the interactions of mutated nsP3 HVD with host G3BP and NAP1 protein families. Thus, in this study, it was used as one of the experimental systems to further dissect the HVD sites that mediate interaction with NAP1 proteins. CHIKV/(3C)/GFP and the designed derivatives, which were used in these and following experiments, also contained GFP gene under control of a viral subgenomic promoter. Its expression simplified evaluation of the spread of the mutants in cultured cells, because some of them were expected to be unable to form distinct plaques.

**FIG. 3.**
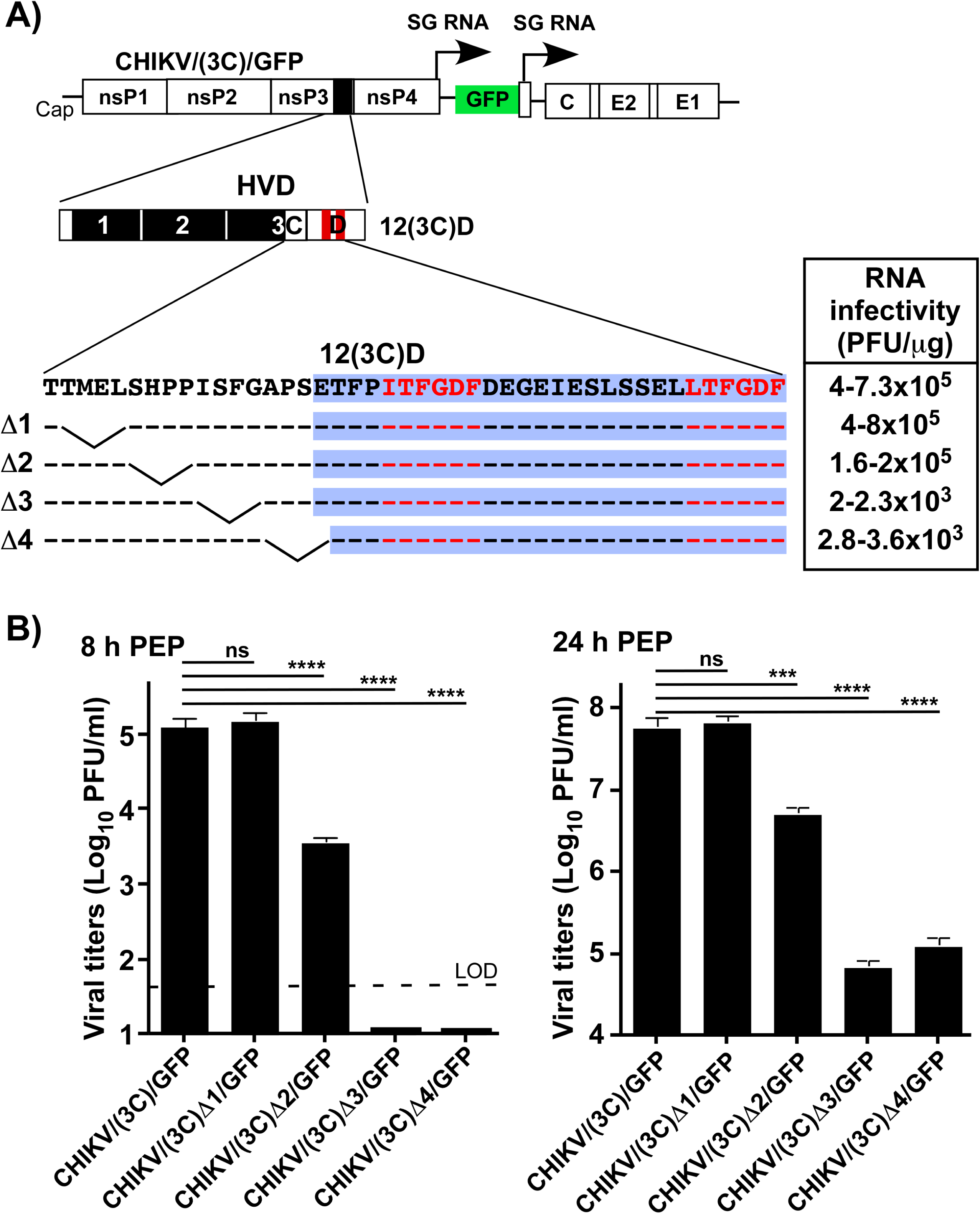
Mutations in NAP1-binding site affect CHIKV replication. (A) The schematic presentation of CHIKV/(3C)/GFP genome, deletions introduced into (3C) fragment and infectivities of the *in vitro*-synthesized RNA in the ICA (see Materials and Methods for details). Dashes indicate aa identical to those in HVD of the parental virus. Fragment D is additionally indicated by blue background. (B) BHK-21 cells were electroporated with 3 μg of the *in vitro*-synthesized RNAs of parental CHIKV/(3C)/GFP and its deletion mutants. Samples of the media were collected at the indicated times post electroporation, and titers were determined by plaque assay on BHK-21 cells. The ICA and titers of the released viruses were assessed in the same experiment, which was reproducibly repeated three times. Means and SDs are indicated. Significance of differences was determined by one-way ANOVA with Tukey test (***, *P***<**0.001; **\*\*\*\***, *P***<**0.0001; n**=**3).

To map the aa in CHIKV HVD, which determine the pro-viral effect of NAP1, we introduced sequential 4-aa-long deletions into (3C) fragment of CHIKV/(3C)/GFP HVD (Fig. 3A). Equal amounts of the *in vitro*-synthesized RNAs were electroporated into BHK-21 cells, and we evaluated effects of the introduced deletions on RNA infectivity using the infectious center assay (ICA) (Fig. 3A), and on the rates of viral replication (Fig. 3B). Deletion Δ1 had no effect on viral replication. Δ2 deletion detectably affected the replication rates, but its effect was not as prominent as those of Δ3 and Δ4. The latter deletions reduced the efficiency of plaque formation in ICA by more than 100-fold (Fig. 3A), and the replication rates of these mutants were a few orders of magnitude lower than that of the parental CHKV/(3C)/GFP (Fig, 3B). A few orders of magnitude lower RNA infectivity was an indication that the detected plaques were likely formed by pseudorevertants, which evolved to a more efficiently replicating phenotype. They could accumulate adaptive mutations either in nsPs or in the *cis*-acting RNA elements of the viral genome. We randomly selected directly in the ICA 4 plaques formed by CHIKV/(3C)Δ3/GFP and CHIKV/(3C)Δ4/GFP. Sequencing the genomes of these plaque-purified viruses revealed the presence of mutations in the structured, N-terminal domains of nsP3. Three isolated pseudorevertants contained a D31A mutation in the macro domain, and another one had a K258R substitution in the AUD. To confirm the positive effects of these mutations on CHIKV replication, they were introduced into cDNAs of the parental CHIKV/(3C)Δ3/GFP and CHIKV/(3C)Δ4/GFP genomes (Figs. 4A and B). The mutations had a profound positive effect on infectivities of the *in vitro*- synthesized RNAs in ICA (Fig. 4B) and increased viral replication rates by two orders of magnitude (Fig. 4C), indicating their true compensatory functions.

**FIG. 4.**
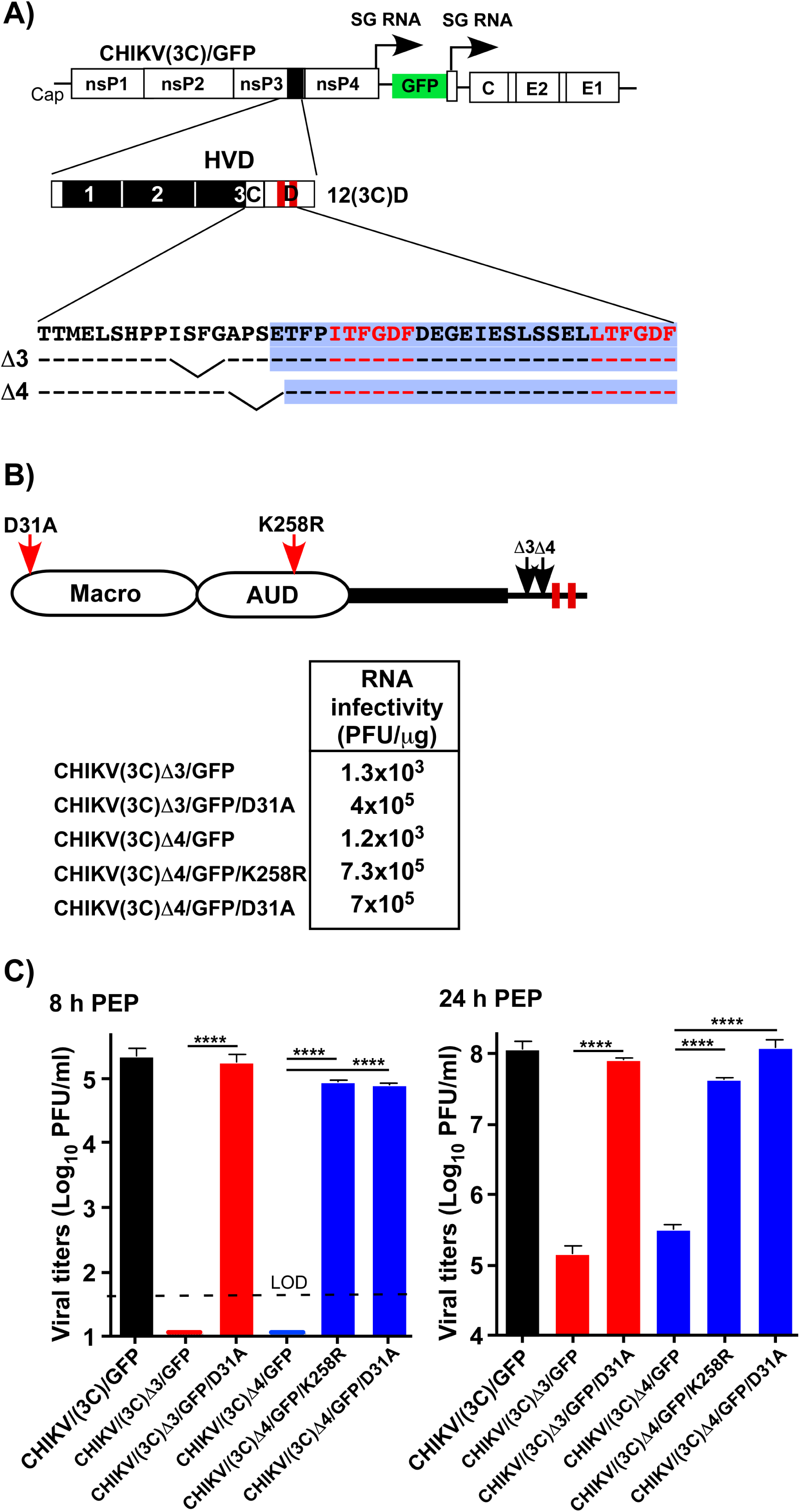
Adaptive mutations in macro domain and AUD compensate negative effects of the deletions in the potential NAP1-binding site. (A) The schematic presentation of CHIKV/(3C)/GFP genome and deletions made in potential NAP1-binding site. (B) The schematic presentation of the domain structure of CHIKV nsP3 in the 12(3C)D variant, positions of the deletions introduced upstream of the G3BP-binding site and adaptive mutations in the macro domain and AUD (indicated by red arrows). Infectivities of the *in vitro*-synthesized RNAs of the parental deletion mutants and those with adaptive mutations in the ICA (see Materials and Methods for details). The experiment was reproducibly repeated twice and the results of one of them are presented. (C) Replication of the variants with adaptive mutations in structured domains and the parental deletion mutants in BHK-21 cells after electroporation of the *in vitro*-synthesized RNAs. Samples of the media were collected at the indicated times post electroporation, and titers were determined by plaque assay on BHK-21 cells. The experiment was reproducibly repeated three times, Means and SDs are indicated. Significance of differences was determined by one-way ANOVA with Tukey test (**\*\*\*\***, *P* **<** 0.0001; n**=**3).

### NAP1L1 interaction with CHIKV HVD is determined by more than one HVD motif

The above data (Figs. 2 and 3) strongly suggested that the binding motif of NAP1L1 is located right upstream of the first G3BP-binding site in CHIKV HVD. To confirm this, we introduced the above deletions Δ1, Δ2, Δ3 and Δ4 into the Flag-GFP-12(3C)D fusion encoded by VEEV replicon (Fig. 5A). The designed replicons were packaged into viral particles and used for infection of the NIH 3T3 cells. As described in the Materials and Methods, the protein complexes were isolated from the cell lysates on anti-Flag MAb magnetic beads and analyzed by Western blot using NAP1L1-specific Abs. Surprisingly, the mutated HVDs were still able to bind NAP1L1 and, as expected, the deletions had no profound effect on the ability of HVD to interact with G3BP (Fig 5B). In contrast, 123D HVD encoding the randomized (3C) peptide was incapable of interacting with NAP1L1 (Fig. 2). CHIKV encoding the latter HVD (CHIKV/123D/GFP) was also nonviable (19). However, smaller modifications, such as 4-aa-long deletions (Fig. 5), did not completely abrogate the ability of CHIKV HVD to co-precipitate NAP1L1. Moreover, the corresponding mutant viruses remained capable of at least very inefficient replication, which ultimately led to their evolution to a more efficiently replicating phenotype (Figs. 3 and 4). Thus, small deletions affected function of NAP1L1 in replication of viral RNA, but did not completely eliminate NAP1L1-HVD interaction. Taken together, these results suggested the possibility that NAP1L1 protein may have an additional binding site in CHIKV HVD. This second interaction could stabilize NAP1L1-HVD complex, but appeared to be insufficient for promoting viral replication in the absence of the wt peptide (3C peptide) upstream of the G3BP-binding site.

**FIG. 5.**
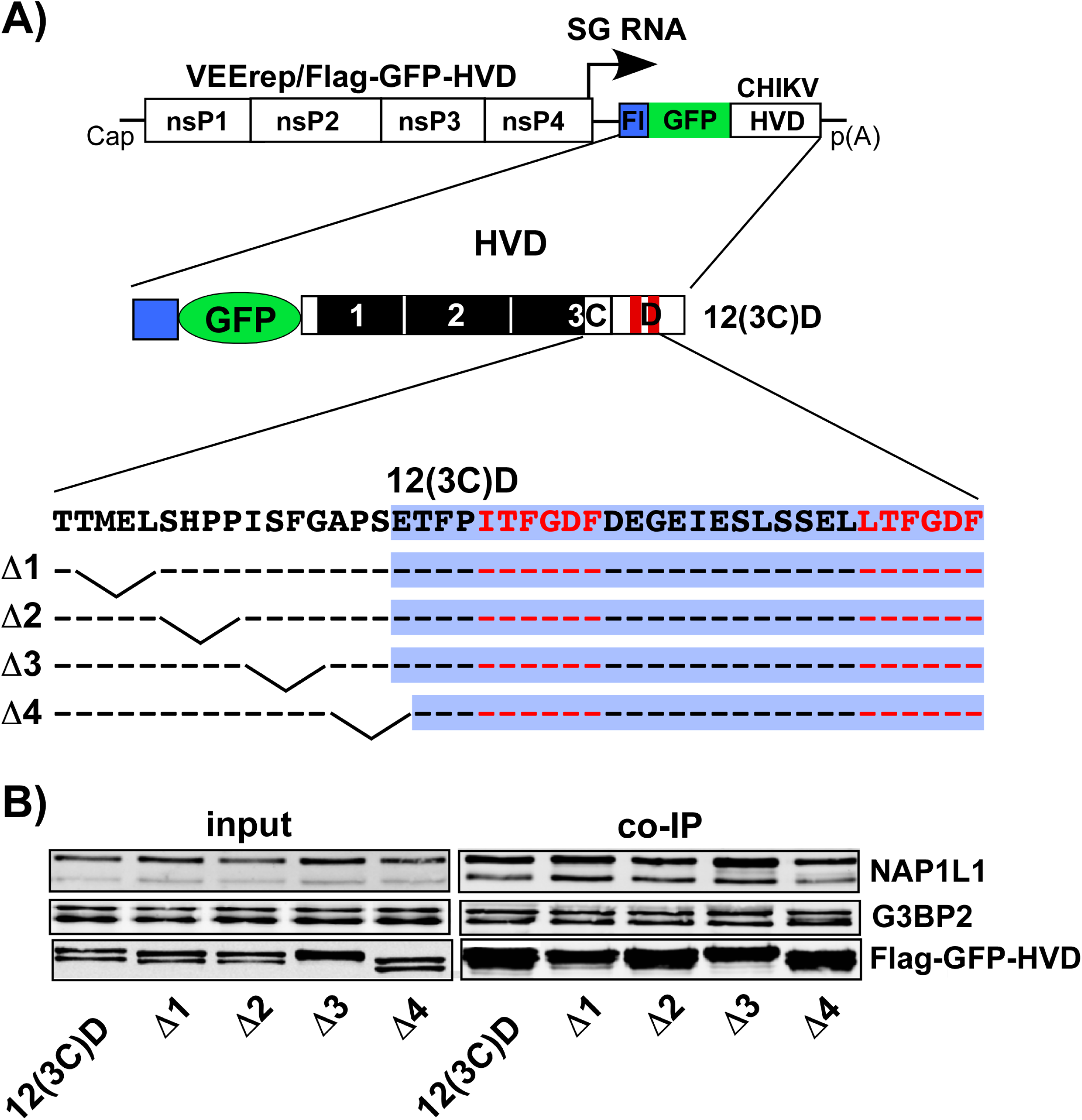
Deletions in putative NAP1-binding site do not abrogate 12(3C)D HVD interaction with NAP1L1. (A) The schematic presentation of VEEV replicon encoding Flag-GFP-12(3C)D HVD fusion with the indicated deletions in the motif located upstream of the G3BP-binding site. (B) NIH 3T3 cells were infected with packaged replicons at an MOI of 20 inf.u/cell. Cells were harvested at ∼3 h p.i. Protein complexes were isolated on magnetic beads as described in Materials and Methods. They were analyzed by Western blot using NAP1L1- and Flag-specific Abs and corresponding secondary Abs. Membranes were scanned on a Li-Cor imager. The experiment was repeated twice with identical results. The results of one of them are presented.

### The C-terminal fragment of CHIKV nsP3 HVD is required for interaction with NAP1L1

To examine the possibility that NAP1L1-HVD interaction is mediated by more than one motif in CHIKV HVD, we designed a set of VEEV replicons encoding Flag-GFP fused with 12(3C)D, which contained additional modifications (Fig. 6A). Since the C-terminal D fragment was the only one left intact in the previous constructs (Fig. 5), we deleted the aa sequences in the very C-terminal peptide located downstream of the first G3BP-binding motif (Fig. 6A). Next, NIH 3T3 cells were infected with these packaged replicons, and the HVD-specific protein complexes were isolated using anti-Flag Mab magnetic beads. Further analyses by Western blot detected no NAP1L1 in the isolated complexes (Fig. 6B), but the mutated HVDs remained capable of interacting with G3BP. This was a strong indication that CHIKV HVD has at least two motifs that determine its interaction with NAP1L1. One of them is located upstream of the first G3BP-binding site and is required for the stimulatory effect of NAP1L1 on viral replication. The second motif is positioned downstream of the G3BP-binding repeat, at the very C-terminus of nsP3 HVD, and its presence alone, without the first upstream peptide, is insufficient for driving CHIKV replication.

**FIG. 6.**
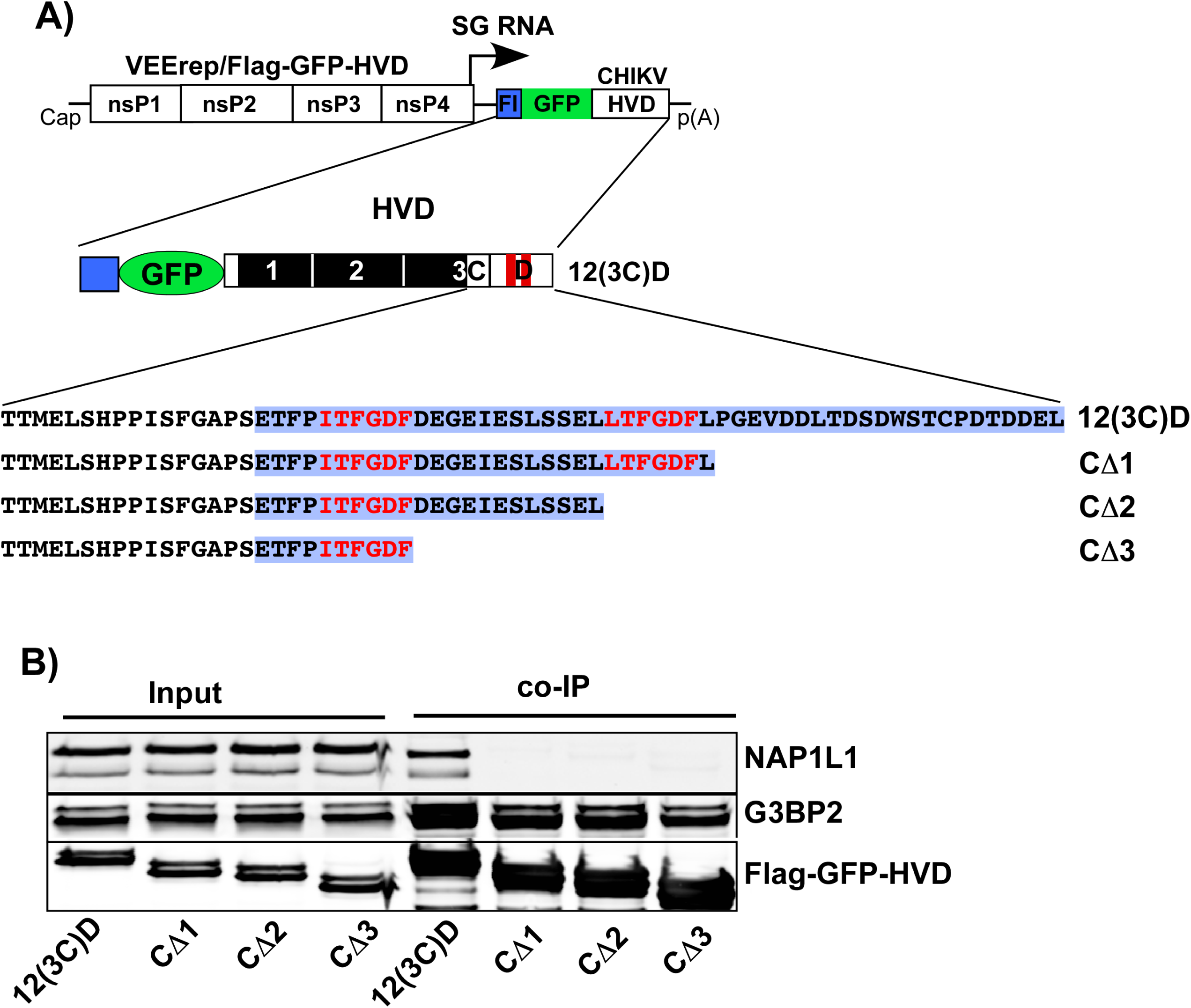
Deletions in the C-terminal fragment of 12(3C)D HVD abrogate its binding of NAP1L1, but not of G3BP. (A) The schematic presentation of VEEV replicons encoding deletion variants of CHIKV 12(3C)D HVD fused with Flag-GFP. Fragment D is additionally indicated by blue background. (B) NIH 3T3 cells were infected with packaged replicons at an MOI of 20 inf.u/cell. Cells were harvested at 3 h p.i. Protein complexes were isolated on magnetic beads and analyzed by Western blot using NAP1L1-, G3BP2- and Flag-specific Abs and corresponding secondary Abs. Membranes were scanned on a Li-Cor imager. The experiment was repeated twice with identical results. The results of one of them are presented.

### Modifications of the C-terminal fragment of CHIKV HVD result in negative effects on viral replication and presence of NAP1L1 in HVD complexes

NAP1 family members are a group of cell proteins that are essential for cell growth, and double knock out (KO) of these genes will likely be lethal for mammalian cells (31–33). Therefore, to demonstrate biological significance of the above finding, we modified the C-terminal fragment of HVD in CHIKV/(3C)/GFP genome (Fig. 7A). The C-terminal fragment was either randomized by changing the positions of the aa or contained a deletion of the 16 aa [CHIKV/(3C)m/GFP and CHIKV/(3C)Δ/GFP, respectively]. The *in vitro*-synthesized viral RNAs were electroporated into the cells, and their infectivities and levels of infectious virus release were evaluated in the ICA and by assessing titers in samples harvested at 24 h post transfection. Modifications introduced into the C-terminus of HVD had deleterious effects on viral replication (Fig. 7A). The latter variants became almost nonviable. Synthesized RNAs exhibited very low efficiencies in the ICA with only pinpoint foci, but not plaques, being detected. Infection spread was almost undetectable and most of electroporated cells remained GFP-negative after 2 days post electroporation. A few orders of magnitude decrease in RNA infectivity indicated that even the detected pinpoint GFP-positive foci were most likely the result of viral evolution and accumulation of adaptive mutations. However, since the infectious titers were very low, this adaptation was not further investigated. Taken together, these data supported the hypothesis that the C-terminus of CHIKV nsP3 HVD is required for the pro-viral function of NAP1 family members.

**FIG. 7.**
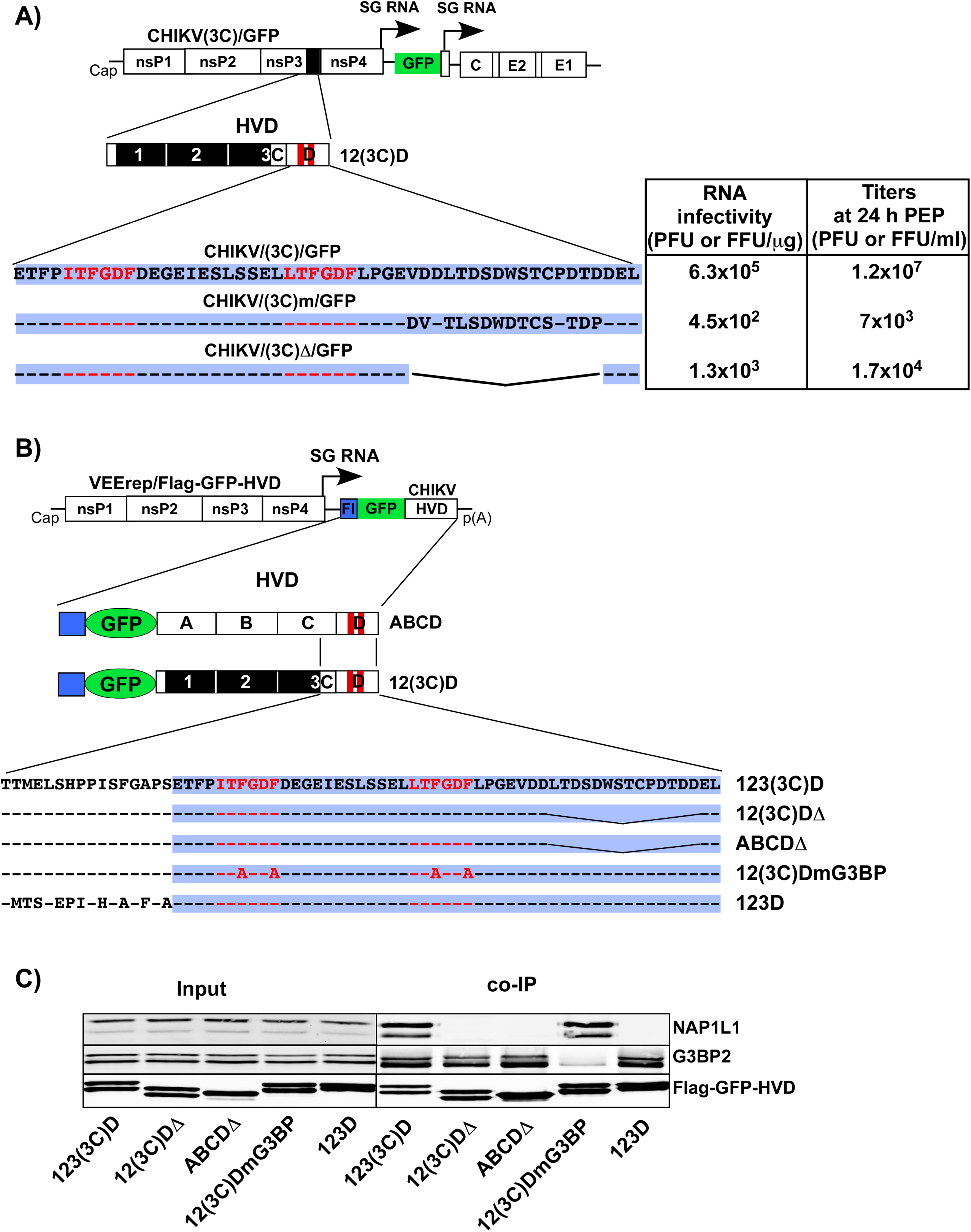
Modifications in the C-terminal fragment CHIKV/(3C)/GFP have deleterious effects on viral replication. (A) The schematic presentation of the genomes and HVDs of the designed viruses, and aa sequences of their D fragments. Dashes indicate aa identical to those in the wt sequence. Table presents infectivities of the *in vitro*-synthesized RNAs in the ICA and viral titers at 24 h post electroporation. RNA infectivities and titers were measured in PFU and GFP-positive foci-forming units for CHIKV/(3C)/GFP and its derivatives, respectively. The experiment was repeated twice with reproducible results. The data from one of them are presented. (B) The schematic presentation of VEEV replicons encoding different variants of CHIKV HVD fused with Flag-GFP. Fragment D is additionally indicated by blue background. (C) NIH 3T3 cells were infected with packaged replicons at an MOI of 20 inf.u/cell. Cells were harvested at 3 h p.i. Protein complexes were isolated on magnetic beads and analyzed by Western blot using NAP1L1-, G3BP2- and Flag-specific Abs and corresponding secondary Abs. Membranes were scanned on a Li-Cor imager.

To experimentally support these data, we expressed additional mutated HVDs as Flag-GFP fusions from VEEV replicon (Fig. 7B). As in the above experiments, HVD complexes were isolated from the NIH 3T3 cells infected with packaged replicons and analyzed by Western blot using NAP1L1-, G3BP2- and Flag-specific Abs (Fig. 7C). Mutations in G3BP-binding repeat abrogated binding of this protein to 12(3C)DmG3BP, but did not affect co-IP of NAPL1L1. The very low level of residual G3BP was likely a result of its binding to the cryptic site of HVD that we had described in our previous studies (19). The deletions of the C-terminal aa sequences in both 12(3C)D and natural ABCD HVDs [12(3C)DΔ and ABCDΔ, respectively] reduced the presence of NAP1L1 in the isolated complexes to undetectable levels, but did not noticeably affect the presence of G3BP. Thus, the results of the co-IP experiments correlated with the inability of CHIKV/(3C)Δ/GFP and CHIKV/(3C)m/GFP mutants to develop spreading infection even in BHK-21 cells (Fig. 7A), which are highly susceptible to CHIKV.

### NAP1L1 is one of host factors having additive stimulatory effects on CHIKV replication

In the next set of experiments, we compared replication of additional CHIKV variants with modified nsP3 HVD (Fig. 8A). One of them, CHIKV/mNAP/GFP, had the NAP1-binding site randomized and thus, was dependent on interactions with all of the CHIKV HVD-specific host factors, except NAP1 family members. Replication of the second one, CHIKV/(3C)/GFP, depended only on binding of G3BPs and NAP1 proteins. Both these mutants and the parental CHIKV/GFP were rescued in BHK-21 cells from *in vitro*-synthesized RNAs. Then human and mouse fibroblasts (MRC-5 and NIH 3T3 cells, respectively) were infected at the same MOI. The modifications introduced into the NAP1-binding site had a negative effect on replication of CHIKV/mNAP1/GFP in both cell lines, which was better detectable at early time p.i. Interestingly, the CHIKV/(3C)/GFP mutant replicated less efficiently in MRC-5 cells than in the NIH 3T3 cell line (Fig. 8B). This was an additional indication that HVD-binding factors function in species- or cell-specific modes, with NAP1 proteins playing a more significant role in CHIKV replication in mouse than in human cells.

**FIG. 8.**
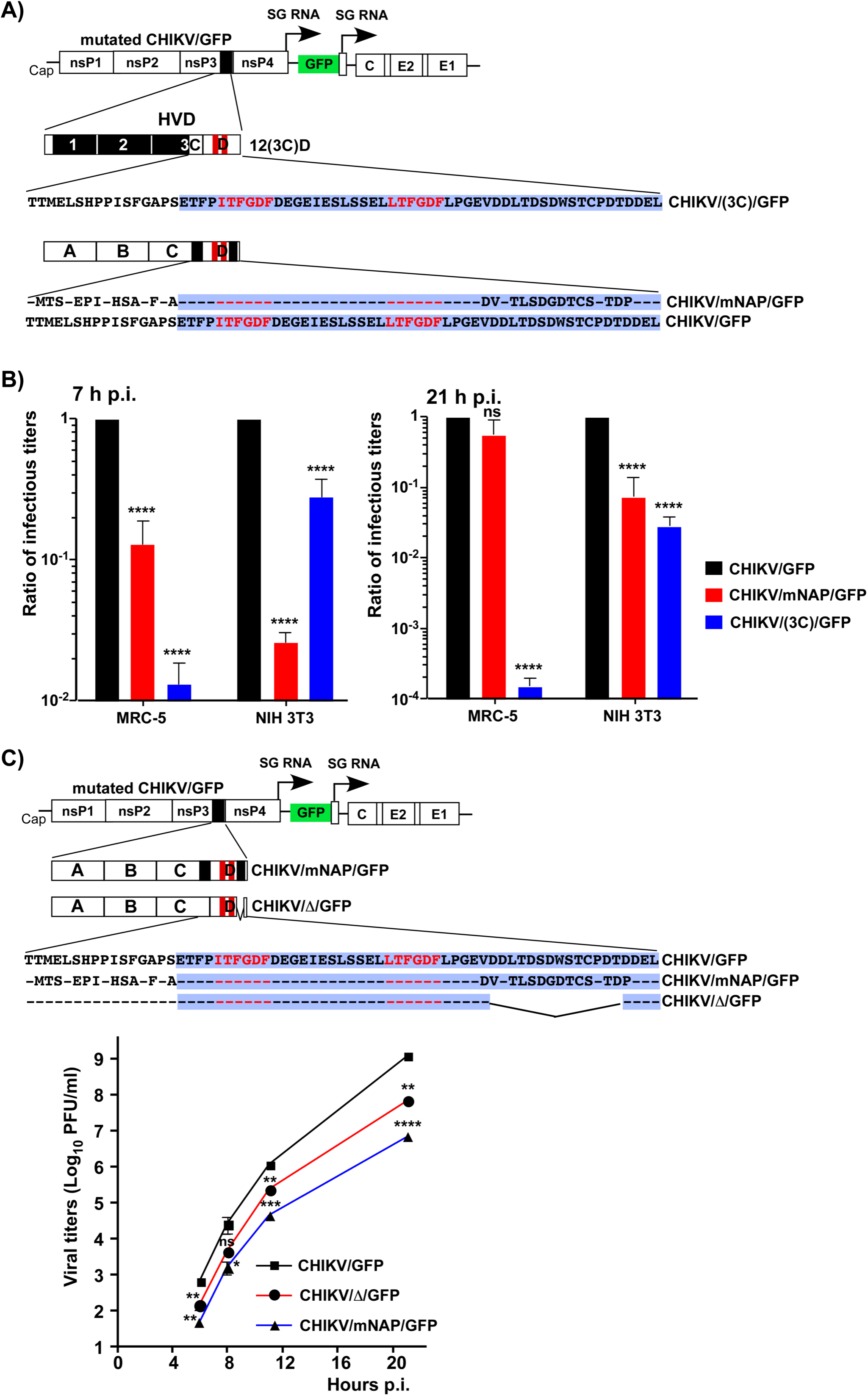
Interaction of CHIKV HVD with NAP1 stimulates viral replication. (A) The schematic presentation of CHIKV/GFP genome, its HVD and the HVDs of the designed mutants, and corresponding aa sequences. Dashes indicate aa identical to those in the wt sequence. Sequence of fragment D is additionally indicated by blue background. (B) NIH 3T3 and MRC-5 cells were infected with the indicated viruses at an MOI of 0.1 PFU/cell, and media were harvested at the indicated times p.i. Infectious titers were determined by plaque assay on BHK-21 cells. Titers were normalized to those of the parental CHIKV/GFP. The experiment was repeated 4 times; means and SDs are indicated. Significance of differences was determined by two-way ANOVA with Fisher LSD test (**\*\*\*\***, *P* **<** 0.0001; n**=**3). (C) The schematic presentation of mutated HVDs in the designed CHIKV variants, and corresponding aa sequences of the modified fragments. NIH 3T3 cells were infected with the indicated mutants and the parental CHIKV/GFP at an MOI of 0.01 PFU/cell. Media were collected at the indicated times p.i., and infectious titers were determined by plaque assay on BHK-21 cells. The experiment was reproducibly repeated 3 times, means and SDs are indicated. Significances of the differences were determined by two-way ANOVA with Fisher LSD test (**\***, *P* **<** 0.05; **, P < 0.01; **\*\*\***, *P* **<** 0.001; **\*\*\*\***, *P* **<** 0.0001; n**=**3).

Next, NIH 3T3 cells were infected with the rescued viruses, which encoded HVD with either the C-terminal NAP1-interacting motif deleted (CHIKV/Δ/GFP) or both motifs located upstream and downstream of G3BP-binding sites were randomized (CHIKV/mNAP/GFP) (Fig. 8C). The rest of their HVDs had wt aa sequences. Both mutants demonstrated slower growth rates in the NIH 3T3 cells (Fig. 8C), but the negative effect of the mutations in both NAP1-specific motifs (see CHIKV/mNAP/GFP) was more prominent than that of the deletion of one of them in CHIKV/Δ/GFP. CHIKV/mNAP1/GFP replicated to almost two orders of magnitude lower titers than the parental CHIKV/GFP.

### NAP1L1 binding requires phosphorylation of CHIKV nsP3 HVD

To further understand NAP1 interaction with CHIKV HVD, we produced NAP1L1 and NAP1L4 proteins in *E.coli* and purified them to homogeneity. Surprisingly, in the *in vitro* binding assay, we did not detect any interaction between CHIKV HVD and NAP1 proteins by size exclusion chromatography and by NMR titration (data not shown), despite these proteins being able to form complexes in vertebrate cells (Figs. 1 and 2). However, alphavirus HVDs are known to be highly phosphorylated (11, 13, 34–41), and proteins produced in *E.coli* lack this post translational modification. Moreover, it was previously suggested, that NAP1L1 interaction with HCV NS5 protein is dependent on phosphorylation (42).

We analyzed the potential phosphorylation sites in CHIKV HVD using NetPhos 3.1 prediction server (43). It predicted that 16 serines and 9 threonines can potentially be phosphorylated (Fig. 9A). Interestingly the C-terminal NAP1-binding motif was predicted to contain 5 phosphorylated aa, all of which could potentially be phosphorylated by CK2 kinase. The recombinant CK2α was found to efficiently phosphorylate the recombinant CHIKV HVD *in vitro* (Fig. 9B). Mass spectrometry analysis of the *in vitro* phosphorylated protein suggested that the major phosphorylated species contains 5 phosphates, and a minor one has 6 phosphates (Fig. 9C). In good correlation with these data, ^31^P NMR suggested that both serines and threonines were phosphorylated and that the ratio of pSer to pThr was 2 to 3 (Figs. 9D and E). Proteomics analysis of phosphorylated HVD detected clearly only a single threonine, T412, phosphorylated at 20% level. However, the C-terminal protein fragment, containing the 16- aa-long peptide with the second NAP1-binding site, could not be detected at all. The most plausible explanation for this was that a high degree of phosphorylation was likely the reason that this peptide could not be identified by conventional mass spectrometry. Combined, these data strongly suggested that the second, C-terminal NAP1-binding site contains 5 phosphorylated Ser and Thr.

**FIG. 9.**
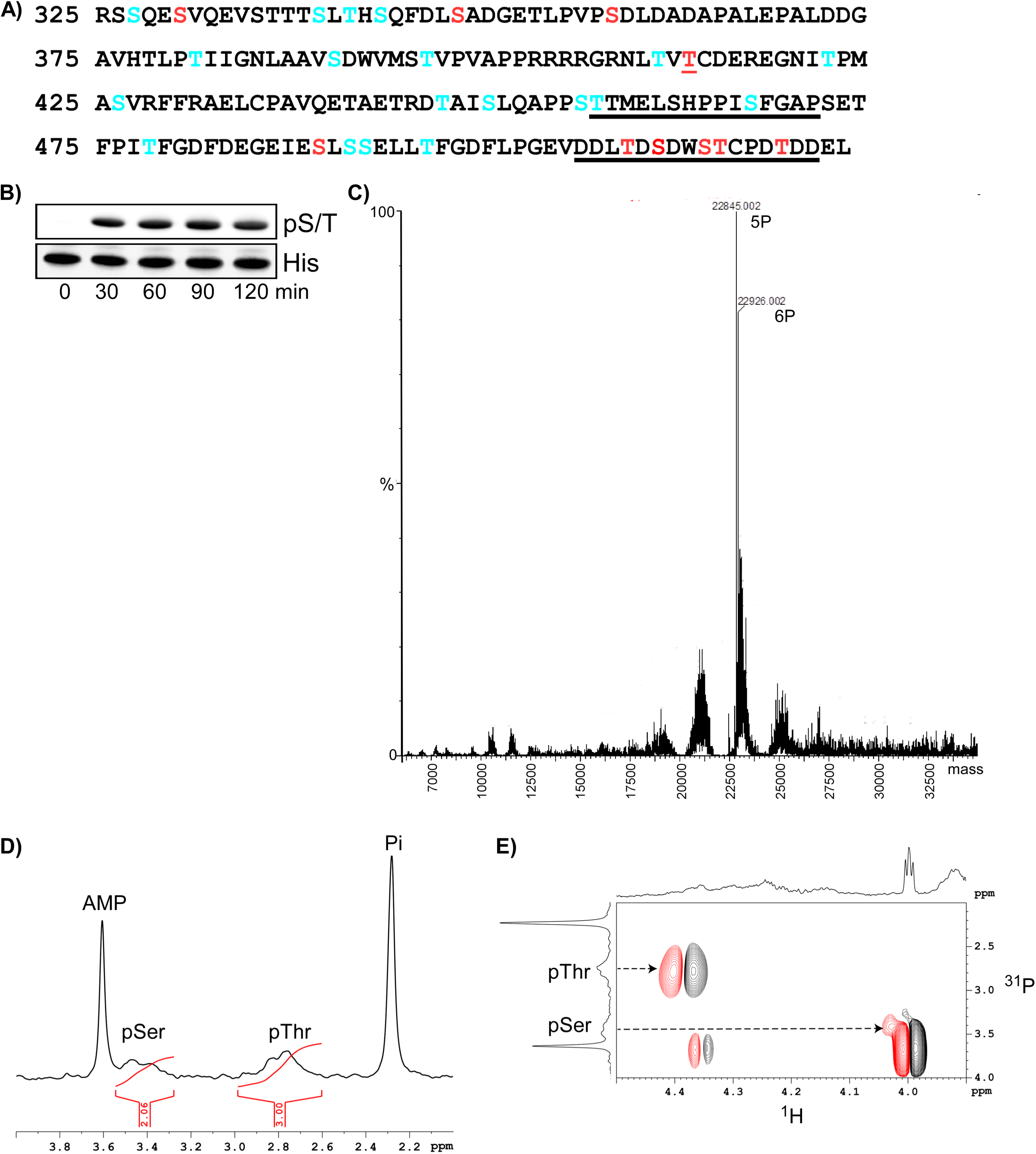
CHIKV HVD is phosphorylated by CKIIα. (A) Potential phosphorylation sites in CHIKV HVD predicted by NetPhos 3.1. The CKIIα-specific sites are indicated in red, other potential sites are indicated in blue. The underlined aa represent the elements of the NAP1-binding sites upstream and downstream of the G3BP-binding motifs. (B) Western blot analysis of *in vitro* phosphorylation of CHIKV HVD-His tag by CKIIα. (C) Charge-deconvoluted spectrum of phosphorylated HVD demonstrates the presence of two major species that contain 5 and 6 phosphates. (D) 1D ^31^P spectrum of the phosphorylated CHIKV HVD (pHVD) between 4.0 and 2.0ppm. Two sharp peaks at 3.61 and 2.29 ppm correspond to AMP and free phosphate, respectively. The broad peaks at 3.48 and 2.76 ppm correspond to phosphorylated serine, labeled as pSer, and phosphorylated threonine, labeled as pThr, respectively. Note that the ratio of the integrals of pSer to pThr is 2 to 3. (E) 2D ^1^H -^31^P correlation spectrum of the pHVD sample is shown. The arrows indicate the connection of the ^31^P resonance of pSer and pThr with corresponding cross peaks.

The *in vitro*-phosphorylated CHIKV HVD was used for the *in vitro*-binding experiments. Individual proteins or their complexes were analyzed by size exclusion chromatography (Fig. 10). HVD was eluted in a single peak. In correlation with the previously published data, both NAP1L1 and NAP1L4 eluted from the column as large aggregates (44, 45). A mixture of NAP1L1 or NAP1L4 with unphosphorylated HVD was eluted as 2 peaks, which corresponded to individual proteins. In the case of phosphorylated HVD, the peak heights for HVD was reduced and formation of a new peak was detected. Gel electrophoresis demonstrated that the latter peak contained both NAP1s and HVD. Interestingly, the HVD/NAP1 complexes eluted later than aggregated NAP1 proteins, which suggested that binding to HVD led to a dissociation of NAP1 aggregates. Thus, nsP3 HVD phosphorylation is a prerequisite of its binding to NAP1 proteins.

**FIG. 10.**
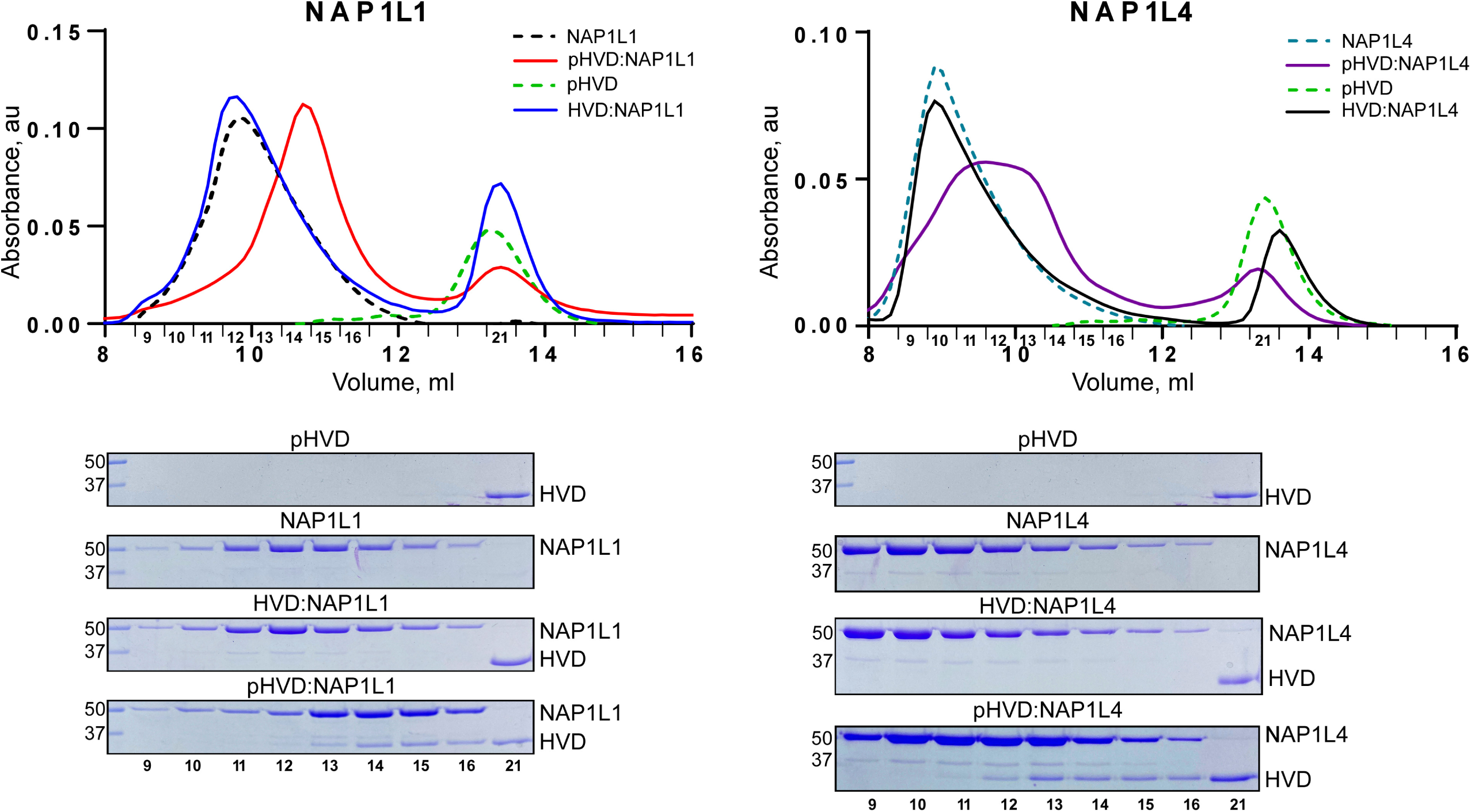
NAP1L1 and NAP1L4 bind *in vitro* only to phosphorylated CHIKV HVD. Analyses of binding were performed by using size exclusion chromatography on Superdex^TM^ 200 increase 10/30 GL column, as described in Materials and Methods. Fractions corresponding to the peaks were analyzed by SDS-PAGE. Gels were stained with Coomassie Blue.

### Inhibition of CK2 kinases prevents NAP1 accumulation in nsP3 complexes and affects viral replication

We next evaluated whether inhibition of CK2 kinase prevents interaction of NAP1 proteins with CHIKV HVD during viral infection. The infected cells were treated with CK2 kinase inhibitor 6-dichloro-1-(β-d-ribofuranosyl) benzimidazole (DRB) and processed for immunostaining with antibodies against nsP3, NAP1L1, NAP1L4 and G3BP1. In mock-treated cells, all three host proteins accumulated in the cytoplasmic nsP3 complexes (Fig. 11). In the presence of DRB, NAP1 proteins no longer co-localized with nsP3. As expected, the DRB treatment did not inhibit interaction between G3BP1 and nsP3 (Fig. 11). This suggested that CK2 kinase is responsible for phosphorylation of CHIKV HVD *in vitro*.

**FIG. 11.**
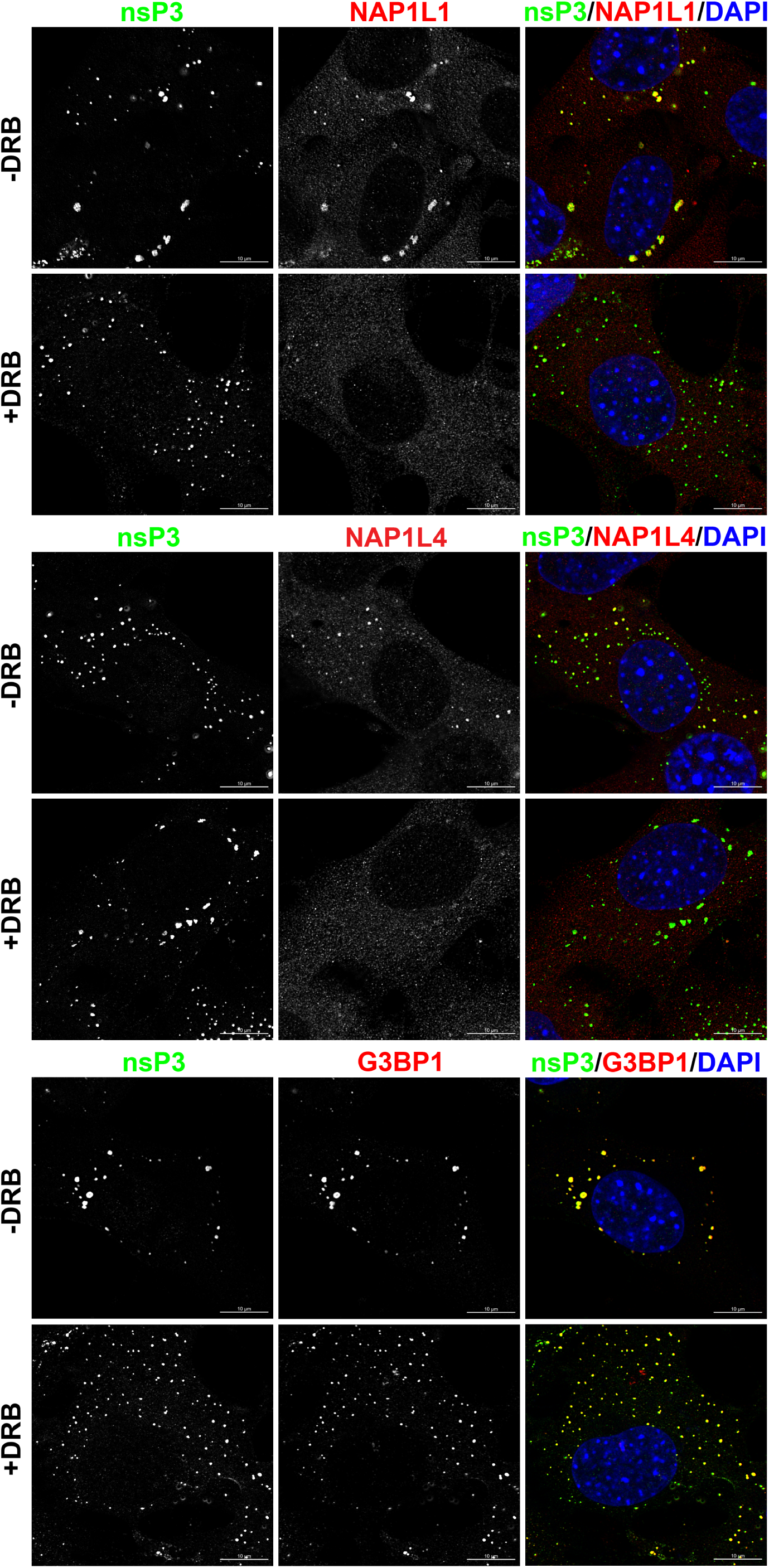
Treatment with DRB prevents accumulation of NAP1 proteins, but not G3BPs, in the cytoplasmic nsP3 complexes. NIH 3T3 cells were infected with CHIKV/GFP at an MOI of 10 PFU/cell and incubated for 15 h in the presence (100 μM) or absence of CK2α inhibitor DRB. After fixing with PFA, cells were stained with antibodies against indicated proteins. Images were acquired on a Zeiss LSM 800 confocal microscope in Airyscan mode with a 63X 1.4NA PlanApochromat oil objective.

In the additional experiments, we evaluated the effect of DRB on CHIKV replication in the NIH 3T3 cells. The negative effect of this drug on viral replication was clearly detectable (Fig. 12). As expected, it was more prominent for CHIKV/(3C)/GFP mutant, whose replication was determined only by G3BPs and NAP1 family members. In case of CHIKV/GFP, other HVD-binding host factors were likely capable of mediating RNA replication, when interaction of HVD with NAP1 was affected by inhibition of the CK2-mediated HVD phosphorylation.

**FIG. 12.**
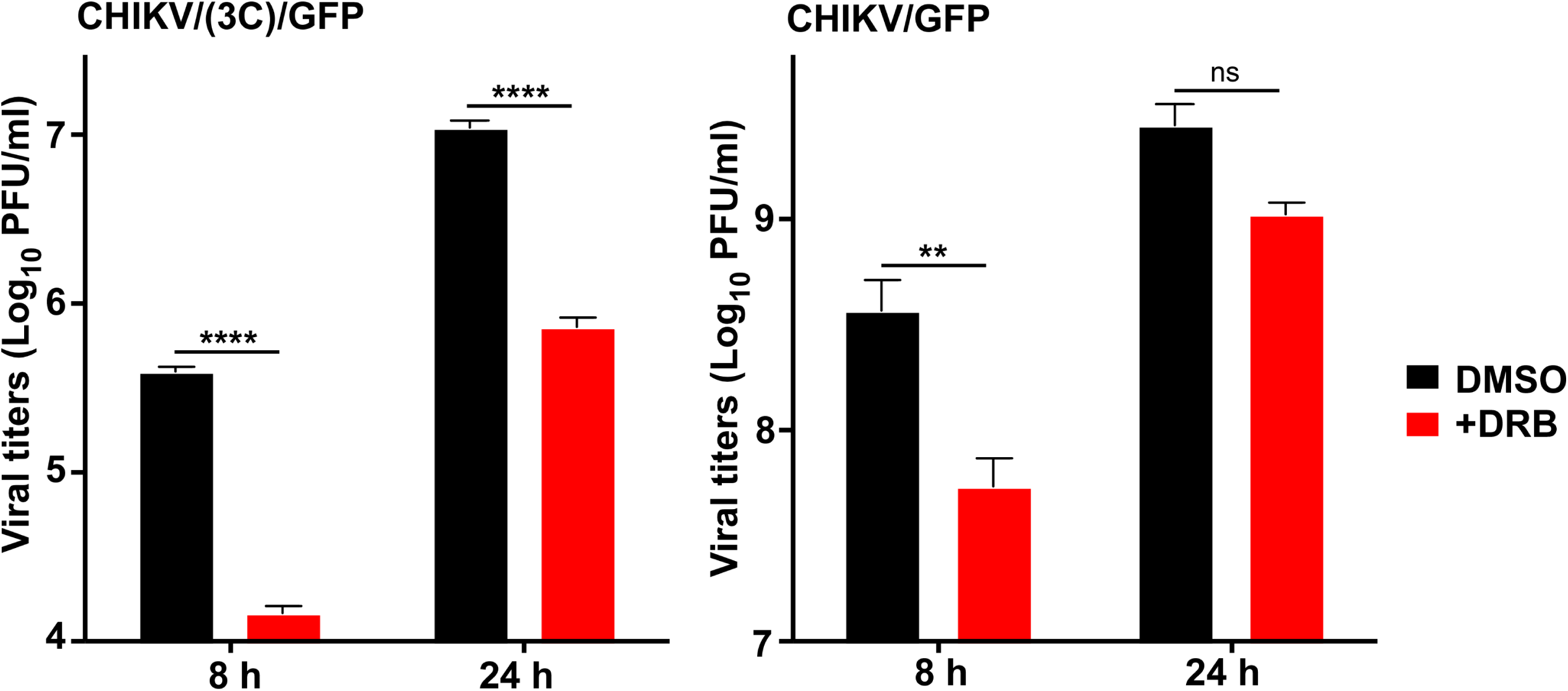
Inhibition of phosphorylation has negative effect on CHIKV replication. NIH 3T3 cells were infected with the indicated viruses in the presence of 100 μM DRB or 0.2% DMSO and after washing with PBS, incubated in the media containing the same concentration of either drug or solvent. Samples of the media were collected at the indicated times p.i., and viral titers were determined by plaque assay on BHK-21 cells. Means and SDs are indicated. Significance of differences was determined by two-way ANOVA with Fisher LSD test (**, P < 0.01; **\*\*\*\***, *P* **<** 0.0001; n**=**3).

## DISCUSSION

Within the last few years, intrinsically disordered proteins (IDPs) and protein domains have attracted a lot of attention, which they deserve. Besides demonstrating unusual physical characteristics, IDPs can interact with multiple proteins, mediate formation of large complexes and thus, are critically involved in numerous cellular processes (46–49). HVDs of CHIKV and VEEV nsP3 and likely nsP3 proteins of other alphaviruses are intrinsically disordered and encode sets of motifs that bind specific host proteins in virus- and cell-specific manners (18, 19, 23, 24, 50–52). These interacting host factors are required for assembly of vRCs, recruitment of G RNA and possibly, for other functions that remain yet to be determined. The disordered structure of HVDs and lack of specific enzymatic activities promote their rapid evolution and adaptation for hijacking different host proteins into alphavirus-specific complexes. Some alphavirus-specific HVDs have even evolved an ability to interact with the same essential host factors by using HVD motifs having different aa sequences (18). For example, the G3BP-binding sites in the HVDs of CHIKV and eastern equine encephalitis virus have different sequences, but both of them recruit G3BP family members to function in viral RNA replication. Similarly, in the HVDs of Venezuelan and eastern equine encephalitis viruses, the critical FXR-binding motifs demonstrate very limited identity, but both function in recruitment of critically important FXR family members (18). The lack of HVD conservation between geographically isolated alphaviruses suggests that these domains may evolve to adapt viruses to mosquito species and vertebrate hosts that circulate in particular areas.

An interesting characteristic of alphavirus HVDs is that in order to support viral RNA synthesis, they can redundantly interact with multiple members of host protein families. In the case of CHIKV, both members of the G3BP family (G3BP1 and G3BP2), three members of the FHL family (FHL1, FHL2 and FHL3) and at least three SH3 domain-containing proteins (CD2AP, BIN1 and SH3KBP1) can function in viral replication with comparable efficiencies (17, 23, 24). The NAP1 family is likely not an exception. Previously, both NAP1L1 and NAP1L4 were readily detectable in co-IP samples of CHIKV HVD-specific complexes (19, 22). Thus, both NAP1L1 and NAP1L4 may redundantly stimulate CHIKV replication.

So far, only the G3BP-HVD interaction was found to be indispensable for CHIKV infection (17). No virus replication or evolution to a more efficiently replicating phenotype was found when both G3BP-binding sites in CHIKV HVD were deleted or mutated or when *G3bp* dKO cells were infected with wt CHIKV. However, CHIKV HVD interaction with G3BPs only is insufficient for viral replication, and other above-described factors demonstrate redundant stimulatory, pro-viral effects. Presence of at least one more interacting motif, besides those specific to G3BP, makes CHIKV viable. Binding of the members of additional families, such as FHL, NAP1 or SH3 domain-containing proteins (BIN1, CD2AP or SH3KBP1), to HVD additively stimulates CHIKV replication. Moreover, the results of our previous NMR-based studies also demonstrated that binding of either CD2AP or FHL1 induces allosteric changes in the distantly located C-terminal HVD peptide, which contains two G3BP-binding motifs (23, 24). These allosteric changes are still difficult to understand, but they may explain the cumulative stimulatory effects of host factor interactions with CHIKV HVD on G3BP-mediated viral replication.

NAP1 family (nucleosome assembly protein 1) contains 5 members both in humans and mice. NAP1L1 and NAP1L4 are the most ancient members and are proposed to form both mono- and heterodimers (28–30, 33). NAP1L1 is likely involved in DNA replication and is detectable in most human tissues and cell lines of human origin with higher expression in rapidly proliferating cells and in tumors. NAP1L1 and NAP1L4 were also proposed to regulate cellular transcription. Both proteins are mostly present in the cytoplasm, but are also detectable in the nucleus. Their shuttling appears to be determined by the encoded nuclear export and nuclear import signals (NES and NLS, respectively) and probably, by their phosphorylation status. The mechanism of NAP1 function in viral replication remains incompletely understood, but CHIKV is not the only virus that hijacks NAP1 proteins for its replication. Previously, NAP1L1 was shown to bind to the disordered fragment of NS5A of HCV, but the biological significance of this interaction remains to be determined (42). In the case of CHIKV infection, NAP1L1/4- HVD interactions have pro-viral effects and likely act directly at the level of viral RNA replication. In our experiments, only fractions of cytoplasmic NAP1L1 and NAP1L4, but not their entire pools, were found in nsP3 complexes (Fig. 1). However, we cannot rule out a possibility that in the infected cells, NAP1 proteins may additionally interact with the fraction of nsP3 that is also diffusely distributed in the cytoplasm outside of the large complexes (53).

An important characteristic of CHIKV HVD, and likely HVDs of other alphaviruses, is that it binds host factors (other than G3BPs and BIN1) with low affinities (21, 23–25). The binding sites are also either relatively long or more than one binding motif is required for interaction with either host protein. The identified FHL1-interacting peptide contains 47 amino acids (23). CD2AP binds simultaneously to two motifs through its SH3-binding domains (24). Moreover, the CD2AP- and FHL1-specific sites overlap, but at least *in vitro*, these proteins are capable of binding to CHIKV HVD simultaneously (23). In this new study, NAP1 function was found to require two HVD motifs, separated by G3BP-binding sites, to express its stimulatory function in viral replication. Moreover, in contrast to other HVD-interacting host factors, NAP1 proteins bind only to the phosphorylated form of the HVD. Phosphorylation of CHIKV HVD by CK2 kinase is required for its binding of NAP1 proteins *in vitro,* and inhibition of CK2 abrogates NAP1/HVD interaction and reduces viral replication. Importantly, it has been demonstrated for Mayaro virus that inhibition of CK2α could also strongly reduce viral replication (54). The requirement for HVD phosphorylation for interaction with other host partners is not so obvious, but might also result in higher affinities of protein binding, and this provides a plausible explanation for the HVD phosphorylation that was described for all known alphaviruses (11, 13, 34–41). Taken together, the accumulating data strongly suggest an additional level of the viral life cycle regulation by phosphorylation of HVD, which leads to differential interactions with host proteins. It would be important to re-evaluate the effect of HVD phosphorylation on interactions with other host factors and identify additional kinases involved in this posttranslational modification of HVD.

The map of interacting motifs in CHIKV HVD that summarizes the currently available data is shown in Fig. 13. It is clear that the C-terminal half of HVD interacts with all of the host factors studied to date. Thus, after binding such a complex combination of cellular proteins, the C-terminal part of CHIKV HVD is likely not as flexible as it is in its free form and might adopt a new conformation. The function of the N-terminal fragment of CHIKV HVD remains unclear. No interacting partners have been determined for this fragment (fragment A), and it can be used for insertion of relatively large proteins, such as GFP or Cherry without strong negative effects on viral replication (16, 55). However, the 62-aa-long deletion in this fragment attenuates CHIKV replication both *in vivo* and *in vitro* (56). To date, the mechanism of attenuation remains unclear. It is possible that either the list of HVD-interacting partners remains incomplete or the distance between structured nsP3 domains and the above described interacting motifs in HVD is important. However, other explanations, such as the essential role of phosphorylation on the latter fragment, are also possible. Taken together, the accumulated data suggest that we are only beginning to understand the complex structure and function(s) of HVD complexes. An additional complicating factor in understanding the HVD function comes from the analysis of adaptive mutations that are selected in response to strong negative effects of HVD-specific mutations on replication of CHIKV and other alphaviruses (13, 16, 51). In this study, the same adaptive mutation, D31A, was detected in in the macro domains of two different CHIKV HVD mutants (Fig. 4). It is in the same position as those we have previously described for the evolved VEEV mutants with strongly modified HVDs (13, 51). Function of the latter mutation in virus replication remains to be determined. However, since it also appears in response to modifications in the promoter elements in the 5’UTR of viral genome (57), it likely represents a universal response to dramatic decreases in viral RNA replication rather than a specific response to modifications in HVDs. In addition, D31 is located in the binding site of ADR-ribose in the alphavirus nsP3 macro domains (14, 58). Mutation of N24A in wt CHIKV nsP3, which inhibit ADR-ribose hydrolase activity of nsP3, requires additional mutation D31G for efficient viral replication (59). Thus, position aa 31 appears to be involved in multiple functions of nsP3.

**FIG. 13.**
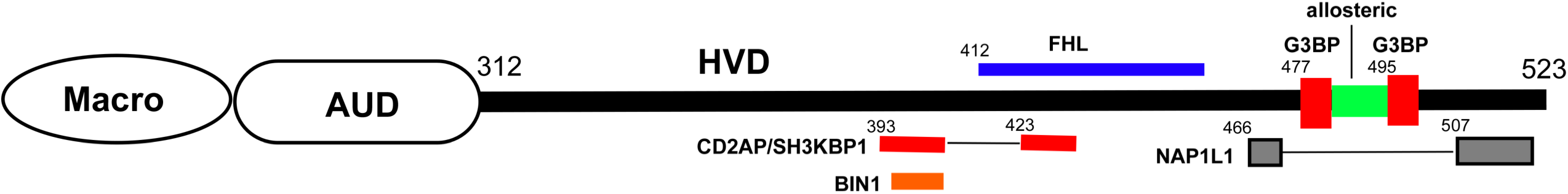
The schematic presentation of the distribution of the binding sites of cellular factors on CHIKV HVD. Numbers indicate positions of aa in CHIKV nsP3.

An additional adaptive mutation was identified in another structured domain of nsP3, the AUD (Fig. 4). Functions of AUD are even less understood, except that introduced mutations may affect RNA synthesis from the SG promoter (60). Both the macro domain- and AUD-specific mutations demonstrate plasticity of alphaviruses and their rapid evolution to higher replication rates and infection spread. However, they likely are of no benefit for the wt virus replication, which has already been optimized by previous evolution.

Taken together, our data demonstrate that i) NAP1-HVD interaction has a strong stimulatory effect on CHIKV replication; ii) both NAP1L1 and NAP1L4 interact with CHIKV HVD and likely have redundant functions; iii) NAP1 family members interact with two peptides, which are located upstream and downstream of the G3BP-binding site of CHIKV HVD, but the NAP1-HVD interaction is independent of G3BP; iv) NAP1 family members interact only with a phosphorylated form of CHIKV HVD, and this phosphorylation is mediated by CK2 kinase; v) inhibitors of CK2 kinase could be explored as a therapeutic means against CHIK infection; vi) NAP1 and other families of host factors have additive stimulatory effects on CHIKV replication.

## MATERIALS AND METHODS

### Cell cultures

The BHK-21 cells were kindly provided by Paul Olivo (Washington University, St. Louis, MO). The NIH 3T3 and MRC-5 cells were obtained from the American Tissue Culture Collection (Manassas, VA). MRC-5 cells were maintained in Dulbecco’s modified Eagle medium supplemented with 10% fetal bovine serum (FBS). Other cell lines were maintained in alpha minimum essential medium supplemented with 10% FBS and vitamins.

### Plasmid constructs

Plasmids encoding infectious cDNA of CHIKV 181/25 containing GFP under control of viral subgenomic promoter (CHIKV/GFP) and its CHIKV/(3C)/GFP derivative have been described elsewhere (61). Their HVD-coding sequences were modified by using gene blocks from Integrated DNA Technologies (IDT) to replace the original nucleotide sequences. The presence of correct sets of mutations was verified by sequencing HVD-coding sequences in the final constructs. Plasmids encoding VEEV replicons with mutated CHIKV HVDs fused with Flag-GFP under control of the subgenomic promoter have been previously described (17).

### Rescuing of the viruses

Plasmids encoding cDNAs of viral genomes with either wt or mutated HVDs (see the figures for details) were purified by ultracentrifugation in CsCl gradients. DNAs were linearized using Not I restriction site located downstream of the poly(A) tail. RNAs were synthesized *in vitro* by SP6 RNA polymerase (New England Biolabs) in the presence of a cap analog (New England Biolabs) in the conditions recommended by the manufacturer. The qualities of the RNAs were evaluated by electrophoresis in nondenaturing agarose gels, and equal amounts of RNAs were used for electroporation without additional purification. Electroporations of BHK-21 cells by the synthesized RNAs were performed under previously described conditions (62, 63). Viruses were harvested at 24 h post electroporation, and titers were determined by a plaque assay on BHK-21 cells (64). To rule out a possibility that the designed CHIKV/GFP mutants require adaptive mutations for their viability, we used an infectious center assay (ICA) to evaluate RNA infectivity. The equal numbers of electroporated cells were seeded on the subconfluent monolayers of naive BHK-21 cells in 6-well Costar plates. After cell attachment, monolayers were covered by 0.5% agarose, supplemented with DMEM and 3% FBS. After ∼60 h of incubation at 37°C, cells were fixed with 4% paraformaldehyde (PFA), and plaques were stained with crystal violet. In some experiments, the release of infectious virus was analyzed directly after RNA electroporation. Aliquots of the media were harvested at the time points indicated in the figures, and infectious titers were determined by plaque assay on BHK-21 cells.

### Packaging of VEEV replicons

The *in vitro*-synthesized RNAs of VEEV replicons and helper, which encodes VEEV TC-83-specific structural genes under control of the subgenomic promoter, were electroporated into BHK-21 cells as described above for viral genomes. Packaged replicons were harvested at ∼24 h post electroporation. Titers were determined in inf.u per ml by infecting BHK-21 cells in 6-well Costar plates (5×10^5^ per well) with different dilutions of the harvested stocks, and numbers of GFP-positive cells were evaluated at 6 h p.i.

### Viral infections

For analysis of viral replication, equal numbers of cells indicated in the figures were seeded into the wells of the 6-well Costar plates. After 4 h of attachment at 37°C, cells were infected at the indicated MOIs in phosphate buffered saline (PBS) supplemented with 1% FBS. After incubation for 1 h at 37°C, cells were washed with PBS and then incubated in corresponding complete media. At the times indicated in the figures, media were replaced, and viral titers were determined by plaque assay on BHK- 21 cells.

### Immunostaining

Cells were seeded into μ-Slide 8-well plates (Ibidi) (2×10^4^ cells/well) and infected at MOIs indicated in the figure legends. In some experiments infected cells were treated with CK2 inhibitor, DRB. At the indicated times p.i., cells were fixed with 4% PFA, permeabilized and stained with specific primary Abs and corresponding fluorescent secondary Abs. Images were acquired on a Zeiss LSM 800 confocal microscope in Airyscan mode with a 63X 1.4NA PlanApochromat oil objective. Antibodies used: anti-NAP1L1 rabbit polyclonal antibodies (14898-1, Proteintech), anti-NAP1L4 rabbit polyclonal antibodies (16018-1, Proteintech), anti-G3BP1 rabbit polyclonal antibodies (gift from Dr. Richard Lloyd) and anti-CHIKV nsP3 (MAB2.19, custom).

### Immunoprecipitations

NIH 3T3 cells and their *G3bp* dKO derivatives were infected with packaged VEEV replicons expressing Flag-GFP fused with different variants of CHIKV HVD at MOIs of 10 or 20 inf.u/cell. Cells were harvested at 3-4 h p.i., when GFP expression began to be detectable under fluorescence microscope. Protein complexes were isolated from the post nuclear fraction of the NP-40-lysed cells using magnetic beads loaded with Flag-specific monoclonal antibody (MAb) (Sigma, Mo) as described elsewhere (17). Presence of proteins of interest in the isolated complexes was analyzed by Western blot using specific Abs and corresponding secondary Abs labeled with infrared dyes. Membranes were scanned on a Li-Cor imager. Antibodies used: anti-NAP1L1 rabbit polyclonal antibodies (14898-1, Proteintech), anti-NAP1L4 rabbit polyclonal antibodies (16018-1, Proteintech), anti-G3BP2 rabbit polyclonal antibodies (HPA018304, Sigma).

### Purification of recombinant proteins

Nucleotide sequence of CHIKV HVD (aa 325-523) was codon-optimized for expression in *E. coli* and His-tag-coding sequence was added to the N-terminus. The synthetic DNA fragment was obtained from Integrated DNA Technologies. It was cloned into pE-SUMOpro-3 plasmid (LifeSensors Inc.) between Nco I and Xho I restriction sites. Plasmid encoding His-HVDchikv (termed HVD thereafter) was transformed into *E. coli* Lobstr BL21 (DE3) RIL (Kerafast), and protein was produced in the M9 media. The expression was induced by 1 mM IPTG after cells reached the density of ∼2 OD_600_. Then cells continued to grow at 37^0^C for 3-5 h. Freshly prepared or frozen cell pellets were lysed in Emulsiflex B15 (Avestin). The lysates were loaded on HisTrap HP column (GE Healthcare) and after extensive washing the recombinant protein was eluted by imidazole gradient. Fractions containing HVD were combined, diluted to contain 25 mM NaCl and further purified on Resource Q column (GE Healthcare). Size exclusion chromatography on a Superdex 75 10/300 column (GE Healthcare) was used as a final purification step.

Nucleotide sequences of NAP1L1 (aa 69-391 of isoform 1, NP_001317160.1) and NAP1L4 (aa 2-375 of isoform 1, NP_001356304.1) were codon-optimized for expression in *E. coli*. The synthetic DNA fragments were cloned into pE-SUMOpro-3 plasmid (LifeSensors Inc.) between Kif I and Xho I sites. The resulting proteins contained His-tags at the N-termini (MGHHHHHHG for NAP1L1 and MGHHHHHHGSR for NAP1L4). These plasmids were transformed into *E. coli* BL21 Star (DE3) (Thermo Fisher), and proteins were produced in M9 media. Protein expression was induced by 1 mM IPTG after cells reached the density of 1-1.5 OD_600_. Then cells continued to grow at 18^0^C for 18 h. Freshly prepared or frozen pellets were lysed in Emulsiflex B15 (Avestin). The recombinant proteins were first purified on a HisTrap HP column (GE Healthcare), followed by buffer exchange using HiPrep 26/10 desalting column (GE Healthcare).

Proteins’ purities and identities were confirmed by SDS-PAGE and mass spectrometry, respectively. In all proteins, the first methionine was present in less than 50% of purified products. Protein concentrations were determined on 280 nm using extinction coefficients, which were determined by ProteinCalculator v3.4 (http://protcalc.sourceforge.net/).

### Analysis of protein interaction by size exclusion chromatography

All size exclusion chromatography (SEC) experiments were performed on the Superdex 200 increase 10/300GL column (GE Healthcare) in phosphate buffer [50 mM N_2_HPO_4_ pH 6.8, 200 mM NaCl, 1 mM tris(hydroxypropyl)phosphine (TCEP)]. For complex formation samples were mixed in equal molar ratios and incubated 15-20 min at room temperature. The 50 μl aliquots were injected into the SEC column and 400 μl aliquots were collected and analyzed by 14% SDS-PAGE.

### HVD phosphorylation

Samples of purified CHIKV HVD were diluted to 100 μM concentration in the buffer containing 25 mM Tris-HCl pH 7.5, 100 mM NaCl, 10 mM MgCl_2_, 1 mM ATP and 1 mM TCEP. Reactions were initiated by adding the recombinant CKIIα to ∼10 μg/ml. Aliquots were collected at the time points indicated in the figure.

Samples were analyzed by Western blot using Abs specific to His-Tag (#66005-1-Ig, Proteintech) and Phospho-CK2 Substrate [(pS/pT)DXE] MultiMab (#8738, Cell Signaling technology) and corresponding secondary Abs labeled with infrared dyes. Membranes were scanned on a Li-Cor imager. The *in vitro*-phosphorylated CHIKV HVD was also submitted to MSBioworks (MSB-service) for identification of phosphorylation sites.

### Analyses of CHIKV HVD phosphorylation by ^31^P NMR

The 1D ^31^P experiment was performed on a Bruker Avance III spectrometer operating at 600.13MHz for ^1^H and 242.94MHz for ^31^P nuclei equipped with a SMART probe. The spectrum was recorded with acquisition times, AQ, of 1.35s, relaxation delay, D1, of 2s and 20480 scans. ^1^H decoupling during D1 and AQ was applied. The sweep width was 50 ppm with the carrier at 0.0 ppm.

The 2D ^1^H -^31^P correlation spectrum was acquired on a Bruker Avance III spectrometer operating at 600.18MHz for ^1^H and 242.96MHz for ^31^P nuclei equipped with a cryo-enhanced QCI-P probe. The spectrum was recorded with acquisition times of 170.4 ms and 10.1 ms in (*t_2_,t_1_*) in 1024 x 64 complex matrices. The sweep widths were 10 and 26 ppm, with the carriers at 4.68 and 0.0 ppm, respectively for ^1^H and ^31^P. A relaxation delay, D1, of 1.5s was employed, and 256 scans were collected. 0.1mM of DSS was added as a reference for chemical shift. The control ^1^H 1D spectrum was collected with acquisition time, AQ, of 1.70s, relaxation delay, D1, of 1s and 16 scans. The sweep width was 16 ppm with the carrier at 4.68 ppm.

All pulse sequences were taken from the Bruker standard library: zgpg30, zgesgp and na_xhcoctetf3gp for 1D ^31^P, ^1^H and 2D ^1^H -^31^P experiments, respectively. Spectra were recorded at 25°C. All NMR spectra were processed by TopSpin 4.0.6 software.

### Analyses of CHIV HVD phosphorylation by native mass spectrometry

Masses of the intact unphosphorylated and phosphorylated CHIKV HVDs were determined using ESI-ToF mass spectrometry (Waters Synapt G2-S(i). Sixty pmol of the proteins were desalted using a Waters BEH SEC column (200Å, 1.7 µm, 2.1 mm X 150 mm) equilibrated in 0.1% formic acid in H_2_O. Spectra were recorded using MassLynx 4.1 in positive ion resolution mode using glu-fib as a lockmass standard. Masses were determined from the acquired spectra using the Waters MaxEnt1 algorithm of MassLynx.

## ACKNOWLEDGMENTS

This study was supported by Public Health Service grants R01AI133159 and R01AI118867 to E.I.F. and R21AI146969 to I.F and Swedish Foundation for Strategic Research grant ITM17-0218 to P.A.

